# A Senescence-Like State is associated with the Immunosuppressive Phenotype of Tumor-Infiltrating Regulatory T Cells

**DOI:** 10.64898/2026.11.12.766305

**Authors:** Temitope Ogunmola, Myung-Chul Kim, Peiyi Zhang, Chandra K. Maharjan, William C. Theune, Kepeng Wang, Ryan H. Kolb, Guangrong Zheng, Nicholas Borcherding, Weizhou Zhang

**Affiliations:** Department of Pathology, Immunology and Laboratory Medicine, College of Medicine, University of Florida, Gainesville, FL 32610, USA; Clinical Pathology, College of Veterinary Medicine, Kyungpook National University, Daegu 41566, Republic of Korea; Department of Medicinal Chemistry, College of Pharmacy, University of Florida, Gainesville, FL 32610, USA; University of Florida Health Cancer Institute, University of Florida, Gainesville, FL 32610, USA; Department of Immunology, School of Medicine, University of Connecticut Health Center, Farmington, CT, 06030, USA; Department of Pathology and Immunology, Washington University School of Medicine, Saint Louis, MO, USA

**Author notes:** Conflict of Interest: G.Z. and W.Z. are equity holders of Dialectic Therapeutics, a company that is developing BCL-X_L_/BCL-2 PROTACs to treat cancers; G.Z., W.Z., P.Z. have also several patents for developing BCL-X_L_/BCL-2 PROTACs to treat cancers.

## Abstract

Regulatory T (Treg) cells are essential mediators of immune suppression within the tumor microenvironment (TME) and represent a major barrier to effective anti-tumor immunity and to therapeutic efficacy of immune checkpoint inhibitors (ICIs). Cellular senescence is traditionally characterized by stable cell-cycle arrest accompanied by distinct molecular and functional changes, yet whether tumor-infiltrating (TI) Treg cells acquire senescence-associated features and how these features influence their immunosuppressive activity remain poorly understood. Here, we demonstrate that TI-Treg cells exhibit both apoptosis- and senescence-associated phenotype while retaining, and in some cases likely enhancing, their immunosuppressive capacity. Compared with peripheral Treg cells, some TI-Treg cells from solid cancers display elevated canonical senescence-associated markers, including p16 and p21, DNA damage response pathways, senescence-associated secretory phenotype (SASP) and anti-apoptotic BCL-2 family genes, as well as activation of secretion associated β-galactosidase (SA-β-Gal). Though physiological Treg senescence leads to diminished immune suppression, TI-Treg cells paradoxically maintain the expression of key immunosuppressive molecules. These findings identify a previously unrecognized senescence-like state in TI-Treg cells that is functionally distinct from the loss of fitness typically associated with senescence in other cell types or aging-associated Treg senescence. Our study establishes senescence-associated reprogramming as a defining feature of TI-Treg cells and highlights the senescence-like state of Treg cells as a potential therapeutic target for selectively disrupting tumor immune suppression while preserving systemic immune homeostasis.

## Introduction

Tumor-infiltrating (TI)-Treg cells have been known for their strong suppressive capacity^1–3^. Treg cells are a major immune suppressive cell type in the tumor microenvironment (TME) ^4–7^ and have been shown to confer resistance to immune checkpoint inhibitor (ICI) therapy ^8–10^. Treg cells also suppress the priming and activation of effector T (Teff) cells in DLNs via constitutive expression of CTLA-4, competing interleukin 2 (IL-2) receptor and/or other potential mechanisms such as LAG3- or TIGIT-mediated immune suppression. Within the TME, several mechanisms have been proposed for TI-Tregs as summarized in many recent review articles^11^; however, the field is still debating on how TI-Treg cells suppress Teff cells in the TME. TI-Tregs express higher levels of T cell exhaustion signature (e.g., CTLA-4, PD-1, LAG3, TIGIT, and ICOS) and T-cell activation signature (*e.g.*, 4-1BB, OX-40, and GITR) molecules ^12^. This is an active Treg phenotype with suppressive activity ^13^. Our data confirmed a similar state of TI-Tregs in ccRCC ^14, 15^, where we identified CD177 expression on the surface of TI-Treg cells from several solid cancers and that CD177^+^ Treg cells exhibit superior immune suppressive capacity to Teff cells^14^. Interestingly, BCL-X_L_ expression is also highly elevated in TI-Tregs from several solid cancers^1^. BCL-X_L_-degrading PROTACs lead to reduction of TI-Treg cells within the TME, along with a consequent activation of CD8^+^ T cells.^1^

BCL-X_L_ is a classic anti-apoptotic BCL-2 family member. BCL-X_L_ expression is correlated with proliferating cells and/or senescence cells, which counteracts potential stress-induced apoptosis related to proliferation and/or senescence. Cellular senescence is a non-proliferative but viable cellular state with prolonged and mostly irreversible cell-cycle arrest ^16–22^. Senescence is characterized by the overexpression of cell cycle inhibitors (such as *CDKN2A* [p16^INK4a^] and/or *CDKN1A* [p21]) ^17, 23^, altered secretome (senescence associated secretory phenotype, SASP) ^24–26^, and deregulated metabolism ^27^ (including the expression of senescence-associated beta-galactosidase, SA-β-Gal) ^28^. BCL-2 family antiapoptotic pathways are elevated during cellular senescence to counteract elevated BH3 family proteins overexpressed during stress. Inhibitors (Navitoclax/ABT-263, A1331852, A1155463 etc.) or PORTAC (DT-2216, PZ15227 or 753b etc.) of the BCL-2 family proteins have been used as senolytics to deplete senescent cells. Though nothing is known about TI-Treg senescence associated with cancer to the best of our knowledge, aging-induced Treg senescence is an emerging field but with some conflicting results ^13, 29–31^. Single cell RNAseq (scRNAseq) indicates an overall increase of aging-associated Tregs and the expansion of an active Treg subset in lymphoid tissues ^13^, in agreement with earlier studies ^30, 31^. At functional level, however, aging-Tregs exhibit signs of senescence and functional loss ^29, 32^. An aging-associated decline of Treg number was found in acutely injured skeletal muscle of aged mice, accompanied by a significant increase in spleen Treg numbers ^33^. These data indicate a tissue-dependent influence of aging on Treg function. As most studies focus on blood/lymphoid organs ^13, 29–31^, the limited available information warrants a complete study on how senescence influences the function of Tregs.

Our findings identify BCL-X_L_ as a critical survival factor for TI-Treg cells and demonstrate that BCL-X_L_-targeting senolytic PROTACs (DT2216 and PZ15227) selectively reduce TI-Treg cell abundance in the tumor microenvironment, as well as in ex vivo human tumor slice culture models.^1^ Building on these observations, we further characterized the senescence phenotype of TI-Treg cells and found that TI-Treg cells express multiple canonical markers of cellular senescence, including p16, p21, senescence-associated secretory phenotype (SASP) genes, DNA damage and repair pathway signatures, and SA-β-Gal activity. Notably, unlike the physiologically senescent Treg cells described previously,^34^ TI-Treg cells exhibiting these senescence features retain, and even upregulate, the expression of key immunosuppressive and activation-associated Treg markers in the current study. Consistent with this phenotype, these cells display enhanced immunosuppressive function, indicating that TI-Treg cells acquire a senescence-like program while maintaining—and potentially augmenting—their suppressive activity within the TME.

## Results

### Human Cancer TI-Treg cells exhibit various immune suppressive states

We have long-term interest in studying human cancer TI-Treg cells. We composed a pan-cancer TIL atlas uTILity that includes 2.6 million cells in 251 patients of 13 cancer types across 27 studies (Table S1). After carefully calling Treg cells based on the consensus of four voters: Azimuth, Monaco, HPCA and the atlas’s own marker gating and removing doublets, 109,015 Treg cells were identified and subject to the Uniform Manifold Approximation and Projection (UMAP). A total of 32 clusters were identified, 31 of which drew from at least 10 cancer types with no study contributing more than half of the corresponding cluster (Fig. 1a). The exception was cluster 22: 88.7% of it comes from one study and its markers are mitochondrial rRNA and pseudogenes, which is excluded from all following analyses. We did thorough analysis of these clusters based on known gene signatures and pathways, including proliferation/cell cycle, apoptosis, T cell senescence, Treg cell suppression, T cell effectors, T cell co-activation etc (Fig. S1a). Cluster 6 is the proliferative Treg cells that expressed MKI67, TOP2A, STMN1 etc (Fig. S1b), along with many other related signatures related to apoptosis and DNA repair pathways etc. While analyzing signature related to cellular senescence using the SenMayo genes, cluster 18 ranked the highest among all the clusters (Fig. S2a), which showed even more prominent enrichment of T cell specific senescence signatures (Fig. 1b). Slingshot for pseudotime trajectory analysis identified a total 15 lineages (Table S2), among which cluster 18 was located at the end of lineage 7 with co-elevation of senescence related pathways such as SASP_chemokines, SASP_proteases, SASP_cytokines as well as T cell senescence (Fig. 1c). Interestingly, though cluster 18 cells were clearly not proliferative (only cluster 6 is proliferative), the cell_cycle arrest signature didn’t elevate along the lineage 7 trajectory (Fig. 1c). ssGSEA analysis comparing with cluster 18 and all other TI-Tregs was performed, which demonstrated that cluster 18 was enriched in pathways related to: regulatory T cell differentiation, replicative senescence, protein secretion, apoptosis, IL2_STAT5 signaling, and oxidative phosphorylation (Fig. 1d) that are known to be involved in Treg cell signaling and metabolic fitness.

**Figure 1.**
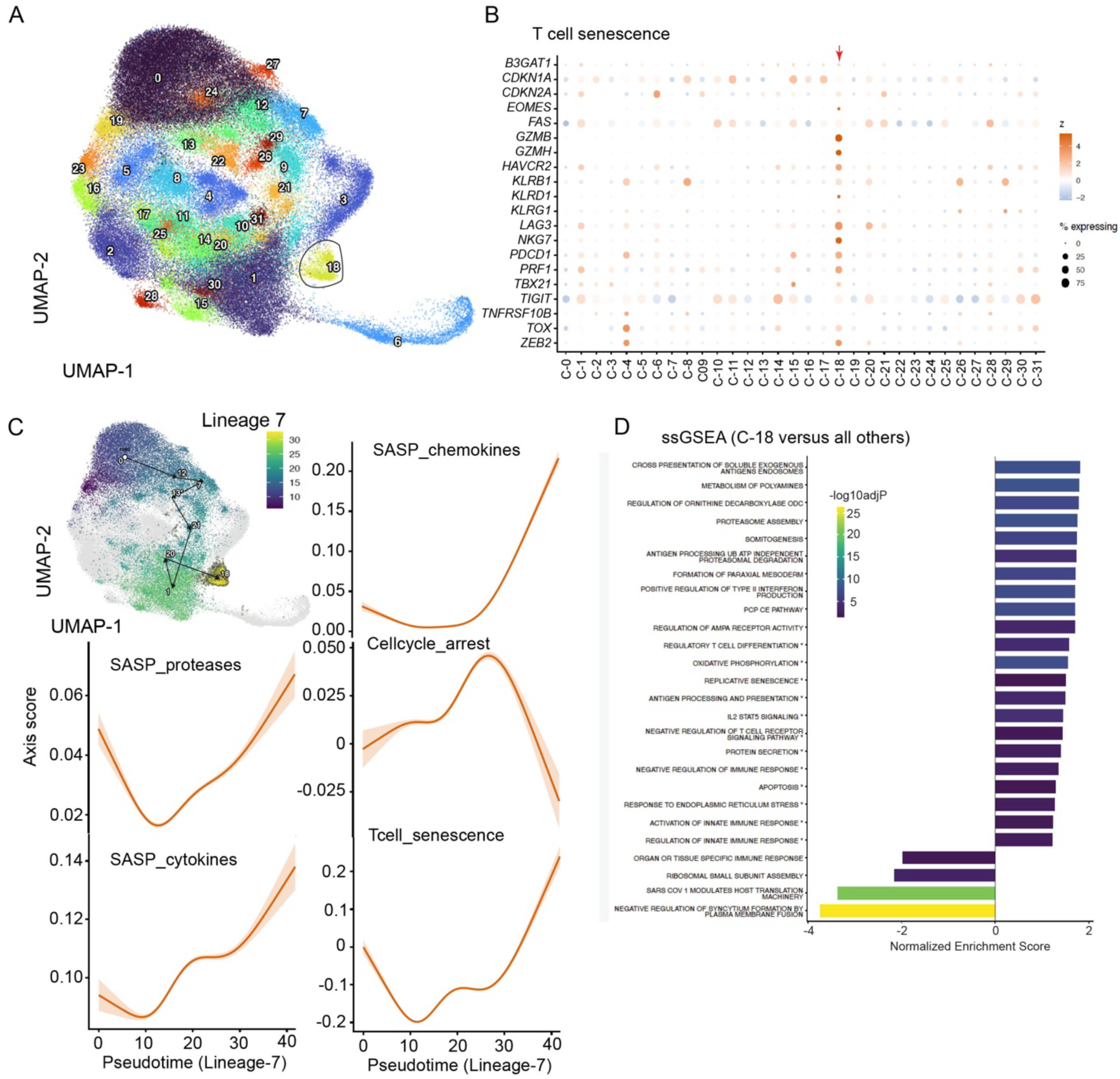
Human Cancer TI-Treg cells exhibit various immune suppressive states. **A-D.** A pan-cancer TIL atlas uTILity was composed including 2.6 million cells in 251 patients of 13 cancer types across 27 studies, within which (A) 109,015 Treg cells were identified and UMAP clusters were shown, noting cluster 18 exhibiting the highest score for T cell senescence. (B) Dotplot showing gene expression patterns related to T cell senescence signatures across all Treg cell clusters. (C) Slingshot for pseudotime trajectory analysis showing lineage 7 starting from root cluster 0 to cluster 18 in UMAP (upper left), along which different senescence-related gene signatures were shown, including SASP_proteases, SASP_cytokines, SASP_chemokines, Cell_cycle_arrest and Tcell_senescence. (D) GSEA analysis showing differential regulated gene sets comparing cluster 18 versus all other clusters. Also see supplementary Figure 1 for additional analysis.

We performed further analysis on T cell senescence signature and identified that cluster 18 exhibited enriched T cell senescence signature genes (Fig. 2a). Cluster 18 had elevated expression of LAG3, IL10, HAVCR2 (TIM-3), PDCD1, TGFB1, and ENTBP1 (CD39) relative to all other clusters combined (Fig. 2b), which were also clearly shown in the expression prevalence scatter plot (Fig. 2c).

**Figure 2.**
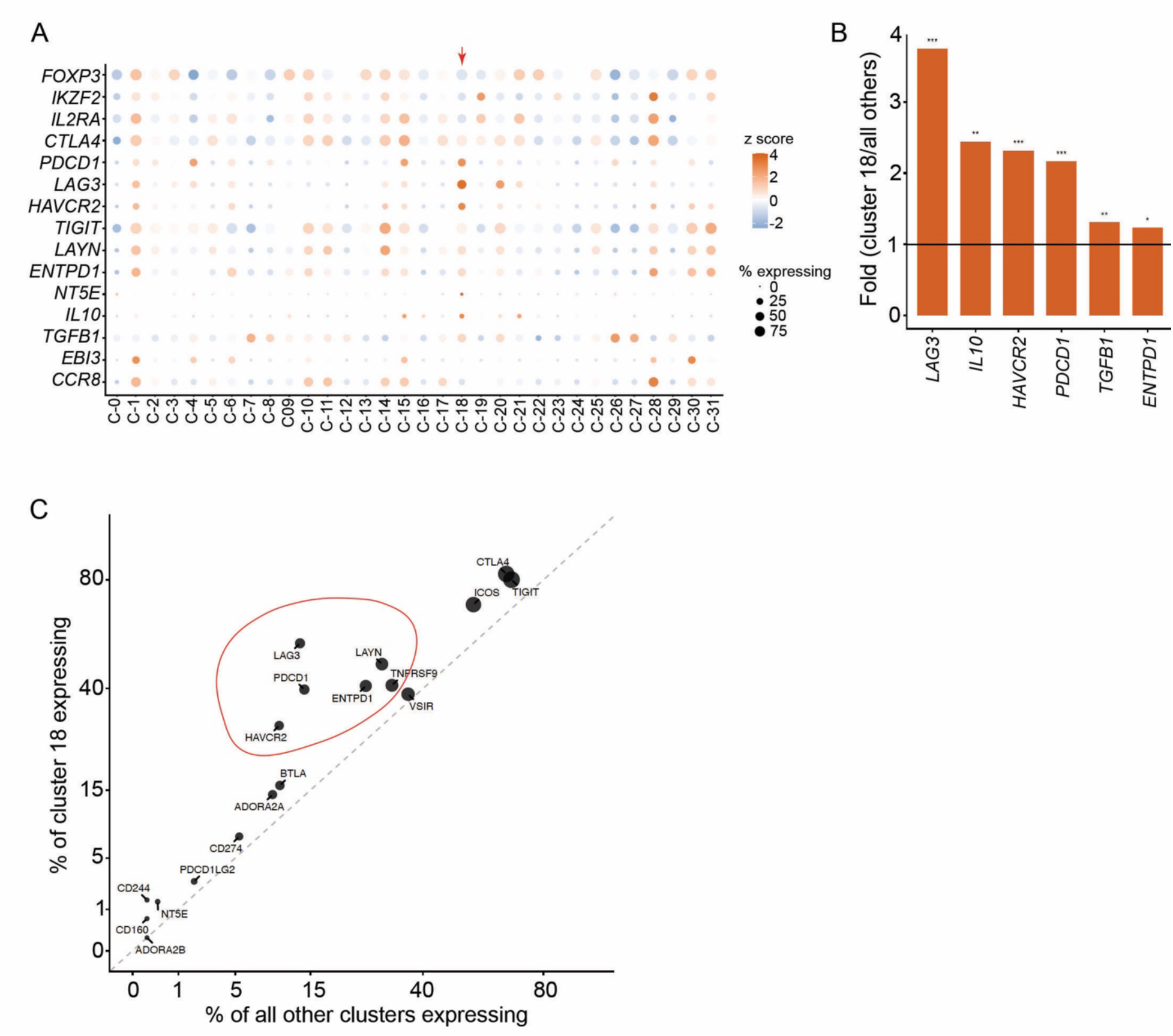
Senescence-like state of TI-Treg cells exhibits distinct set of immune suppressive signature. **A.** Dotplot showing Treg suppressive gene expression patterns related to all Treg cell clusters. **B.** Bar graph showing major Treg suppressive genes enriched in cluster 18 relative to all other clusters. **C.** Percent of cells expressing Treg suppressive genes in cluster 18 relative to all other clusters. Also see supplementary Figure 2-3 for additional analysis.

Aging related Treg cell senescence was recently reported, with a loss of immune suppressive function and believed to be involved in aging associated chronic inflammation. To the best of our knowledge, there is no systematic report related to TI-Treg cell senescence within human cancer. Curiously, we compared SenMayo signatures between paired blood Treg cells and TI-Treg cells, and there was a general trend of elevated SenMayo senescence signature in TI-Treg cells across all tested cancer types including biliary tract cancer, colorectal cancer, head and neck squamous cell carcinoma, lung cancer, melanoma and renal cancer (Fig. S2b).

In addition, we also re-analyzed our published bulk RNAseq datasets from MC38 tumor-bearing mice by comparing Tregs from spleen (SP-Tregs) and those from tumors (TI-Tregs) (Fig. S2c). DEGs that are upregulated in TI-Tregs include—1) core senescence signature genes such as Cdkn2a, Cdkn1a, Glb1, SASPs (Cxcl2, Mmp9, Mmp12, Serpine1, Igfbp7, Tgfb1, Tnf, Il1a, Il1b, Cxcl10, etc.); 2) apoptotic signature genes: co-elevation of both Bcl2l1 and Mcl1 anti-apoptotic members as well as Bak1, Bad, Pmaip1, Bok, Casp3 pro-apoptotic genes; 3) Treg suppressive signatures: cytokines (Tgfb1, Il10, Ebi3), immune checkpoints (Tigit, Lag3, Tim-3, Pdcd1), co-activation markers (Icos, Tnfrsf18, Tnfrsf9, Tnfrsf4), effectors (Gzmb, Prdm1), and metabolic modulators (Entpd1/Cd39 and Arg1)—all of which are known to be essential or associated with Treg suppressive functions (Fig. S2c). These TI-Treg signatures from mouse tumors are largely in agreement with those from human cancers.

We have recently reported that TI-Treg cells in RCC are clonally expanded.^1^ Looking into clonality of Treg cells, cluster 18 ranked the 2^nd^ with more than 25% of Treg cells exhibiting certain level of expansion (Fig. S3a). Treg apoptosis has been described as one of the immune suppressive mechanisms in ovarian cancer and hepatocellular carcinoma. We performed correlation study and found that cluster 18 ranked the top in T cell senescence score, but exhibited intermediate level of apoptotic signature (Fig. S3b).

Here we described that human and mouse TI-Treg cells exhibit a senescence-like state; however, these TI-Treg cells share a common elevation of immune suppressive signature including same set of genes (Lag3, Pdcd1, Entpd1, Tgfb1, Il10), suggesting different outcomes of senescence-like state within the TI-Treg cells and aging-related senescent Treg cells.

### TI-Tregs from ccRCC express high levels of senescent markers

Working with human TI-Tregs is relatively challenging due to the limitation of specimen size and low frequency of TI-Tregs. Renal cell cancers (RCC) are known to have larger mass in clinic. We have recently defined CD177^+^ TI-Tregs from human RCCs and other solid cancers^1^ and further analyzed other senescent markers using our recently published scRNAseq dataset (GSE121638)^15^. UMAP clustering identified all major immune cell types (Fig. 3a). Additional analysis of Treg cells formed 4 clusters (Fig. 3b) and the cluster 2 exhibited significant enrichment of senescence gene signature (Fig. 3c). We determined the activity of SA-β-Gal in TI-Tregs from human RCC using a novel tracer NIR-BG2 ^35^ (Fig. 3d). TI-Tregs from human ccRCC exhibit significant SA-β-Gal relative to paired blood Tregs or fluorescence-minus one (FMO) controls (Fig. 3d). TI-Treg cells from ccRCC have elevated levels of *CDKN2A*/p16, *CDKN1A*/p19, and *BCL2L1* (Fig. 3e)^1^; however, TI-CD4^+^ conventional T cells (Tconv), -CD8^+^ T cells, or tumor-associated macrophages (TAM) exhibit varying levels of these genes (Fig. 3e).

**Figure. 3.**
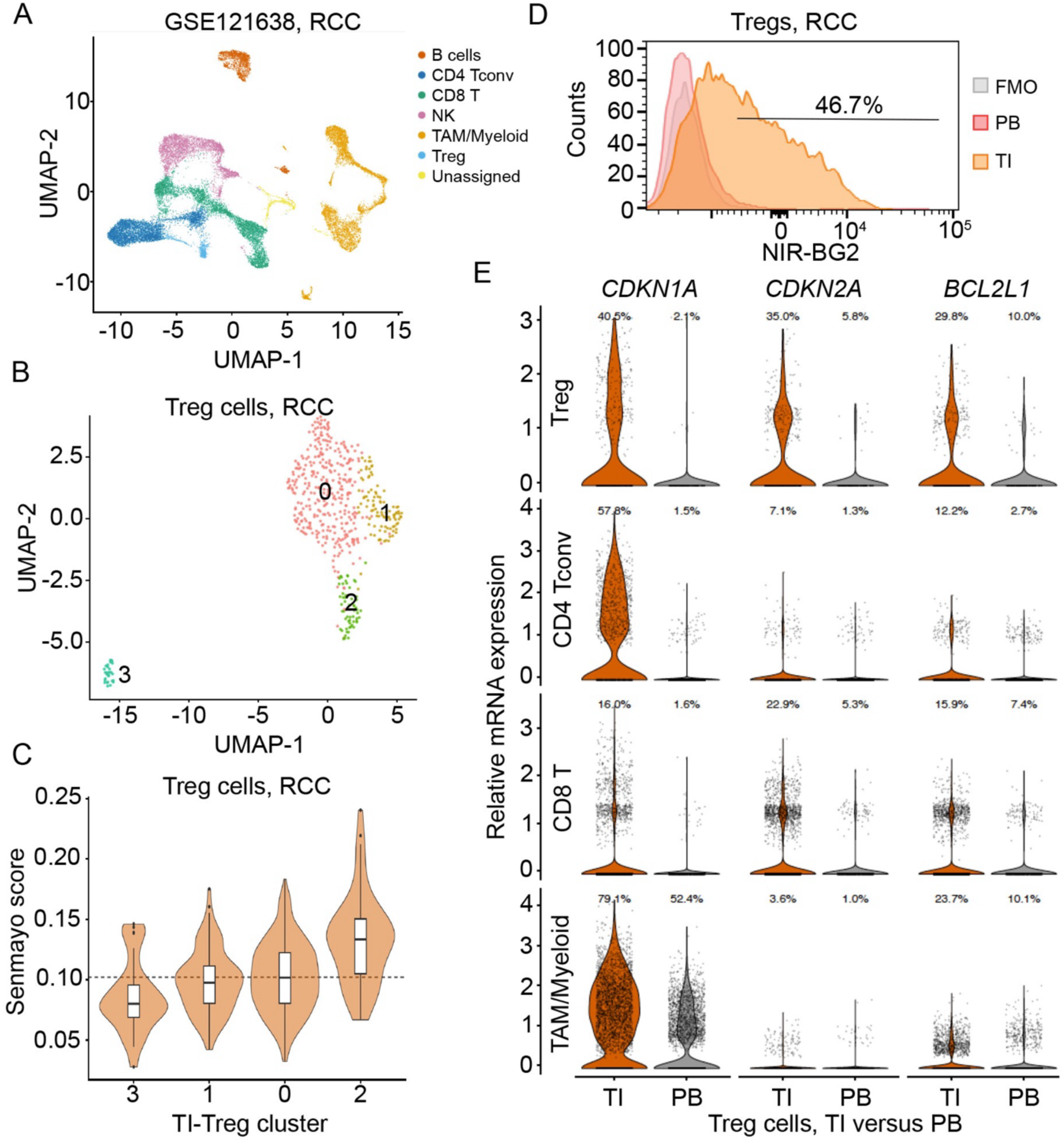
TI-Treg cells from human RCC exhibit senescence signatures. **A.** UMAP clustering of all immune cells from our published GSE121638 single cell dataset, including 3 paired ccRCC specimens and corresponding blood cells. **B.** UMAP analysis of Treg cells within A showing 4 distinct populations. **C**. Violin plot showing that TI-Treg cells exhibited different levels of senescence signature gene set. **D.** Flow cytometric analysis showing increased SA-β-Gal activity in TI-Tregs from human ccRCC patient specimens (n=3 specimens). **E**. Violin plots showing senescence-associated genes in Treg cells, CD4 conventional T cells, CD8 T cells, as well as myeloid cells. TI, tumor infiltrating; PB, peripheral blood.

### Induction of Treg cell senescence and immune suppression by renal cancer cells

Since TI-Treg cells exhibit SA-β-Gal activity and other senescent signatures, we reason that the induction of Treg cell senescence depends on the TME. We first determine whether cancer cells can directly induce Treg cell senescence by co-culturing human blood Treg cells, either with HK-2 — an immortalized human proximal tubule epithelial cells (normal), or renal cancer cells (RCC) (Fig. 4a–4b). Treg cells exhibited much more pronounced SA-β-Gal activity when co-cultured with RCCs than with HK-2 cells (Fig. 4a) as quantitated (Fig. 4b, RCC-Tregs versus normal-Tregs).

**Fig. 4.**
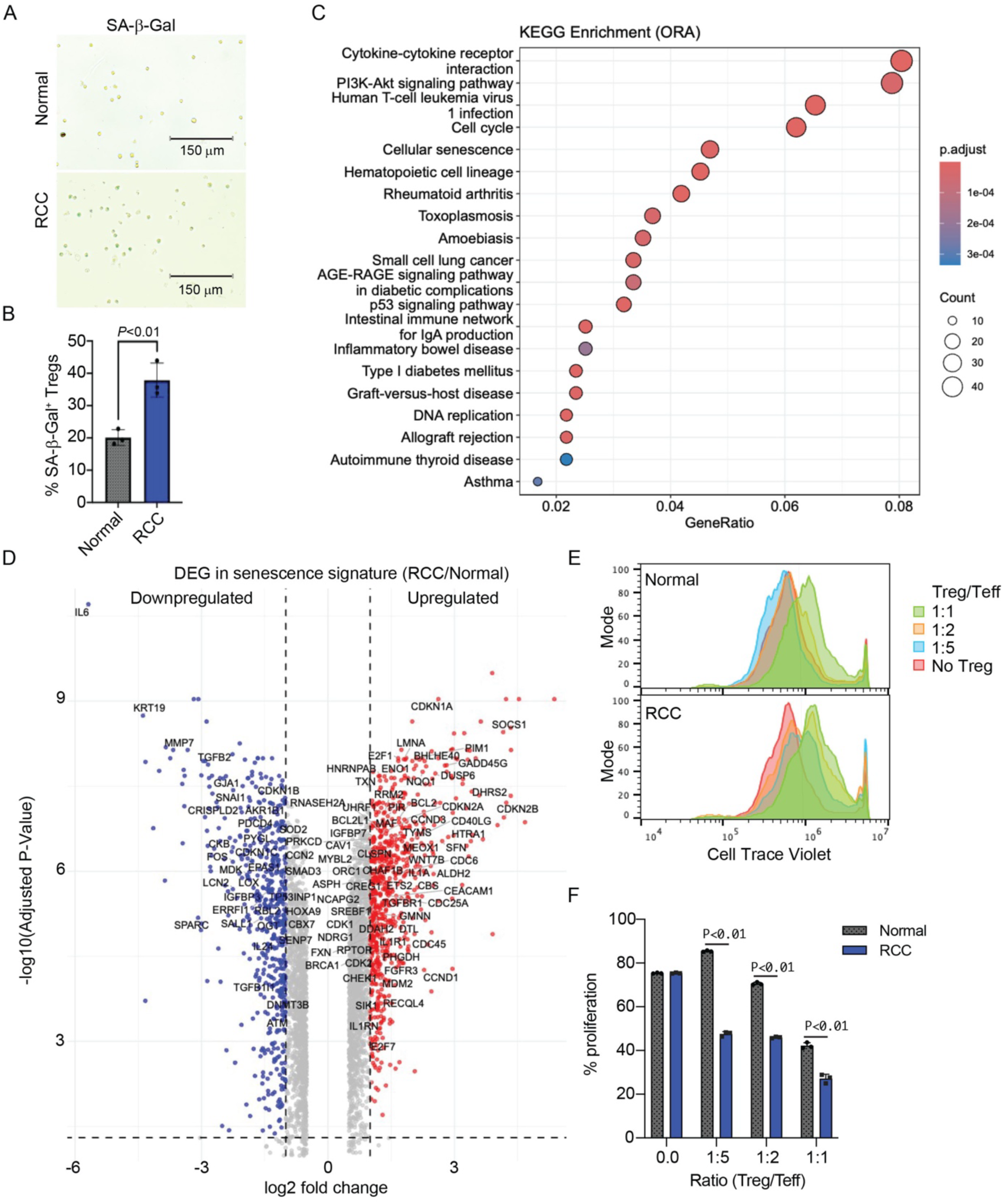
Renal cancer cells induce Treg senescence and enhance Treg suppressive function. **A.** Representative images of SA-β-Gal activity detected by X-gal staining in human Tregs co-cultured for 72 hrs with normal kidney epithelial cells (HK-2) or renal cancer cells (769-P). **B.** Percentage of SA-β-Gal⁺ Tregs ± s.d. under each condition (n=3 biological replicates). Scale bars, 150 μm. **C.** KEGG pathway enrichment analysis depicting the 20 most enriched pathways among genes differentially expressed in RCC relative to Normal co-cultured Tregs by bulk RNA-seq. **D.** Volcano plot depicting the fold difference in expression of senescence signature genes in RCC relative to normal co-cultured Tregs. Dashed lines indicate thresholds of log2 fold change ±1 and adjusted P value of 0.05; grey indicates no significant difference. **E.** Representative histogram showing CellTrace Violet dilution in effector T cells (Teff) cocultured with normal or RCC co-cultured Tregs at the indicated Treg/Teff ratios, and **F.** percentage of proliferating Teff ± s.d. at each ratio (n=3). Unpaired T-tests (B, n=3; and F, n=3 biological replicates, two-sided) were performed and the corresponding P values are indicated. **A, B, E, F.** Normal, Tregs co-cultured with HK-2 normal kidney epithelial cells; RCC, Tregs co-cultured with 769-P renal cancer cells. **C.** Count, number of enriched genes per pathway; GeneRatio, fraction of pathway genes recovered; p.adjust, Benjamini-Hochberg adjusted p-value. **E.** No Tregs, Teff cultured without Tregs as control. Also see supplementary Figure 4 for additional analysis.

At the transcriptional level, we performed bulk RNAseq analysis to compare changes between normal-Treg and RCC-Treg cells. As expected, based on SA-β-Gal staining, we found that RCC-Treg cells exhibit typical pathway changes including cytokine-cytokine receptor interactions (SASP), cellular senescence, p53 signaling pathway, etc. (Fig. 4c). Further breaking down the differentially expressed genes (DEGs) related to senescence, RCC-Treg cells exhibited many core senescence genes such as *CDKN1A*, *CDKN2A*, *BCL2L1*, etc. (Fig. 4d), as well as those genes known to mediate the immune suppressive functions of Tregs, such as *TNFRSF18* (GITR), *TNFRSF4* (OX40), *ENTPD1* (CD39), *BTLA*, *CCR8*, *HAVCR2*, and *IL2RA* (CD25) (Supplementary Fig. S4a).

To test whether these Treg cells are immunosuppressive, CD4^+^ Teff cells from same PBMCs were co-cultured with either Treg-normal cells or Treg-RCC cells from Fig. 4a-4b with different ratios. Treg-RCC cells exhibited stronger suppression to T cell proliferation as shown by flow cytometry (Fig. 4e) and summarized in Fig. 4f, resembling the highly immune suppressive nature of TI-Treg cells.

All above results support the induction of Treg cell senescence by cancer cells within the tumor microenvironment, while enhancing the immune suppressive capacity toward Teff cells. To understand the molecular mechanism of senescence induction of Treg cells, we cultured primary blood Treg cells with different doses of hydrogen peroxide and found that oxidative stress – a known mechanism to induce Treg apoptosis and immune suppression – significantly enhanced senescence measured by SA-β-Gal activation (Fig. S4b)

### ENTPD1 (CD39) is a surrogate marker for senescent TI-Treg cells

Currently there is no well-established surface markers to separate viable senescent Treg cells versus non-senescent Treg cells. We explored different Treg-related surface markers, using NIR-BG-2 to label and distinguish senescence cells from non-senescent cells within PBMC or TI-Treg cells. Using paired human PBMC and RCC specimens, we found that a very small population of KLRG1+CD25+ Treg cells (1.04%) (Fig. 5a) as well as small population of CD39+CD25+ Treg cells (1.63%) (Fig. 5b) in the parental CD4+ T cell gating from human PBMC. Interestingly, CD39+CD25+ Treg cells (Fig. 5c) were partially stained positive for NIR-BG-2, while most KLRG1+CD25+ Treg cells were positive for NIR-BG-2 staining, supporting that KLRG1+ is a potential surrogate marker for senescent Treg cells in PBMC.

**Figure 5.**
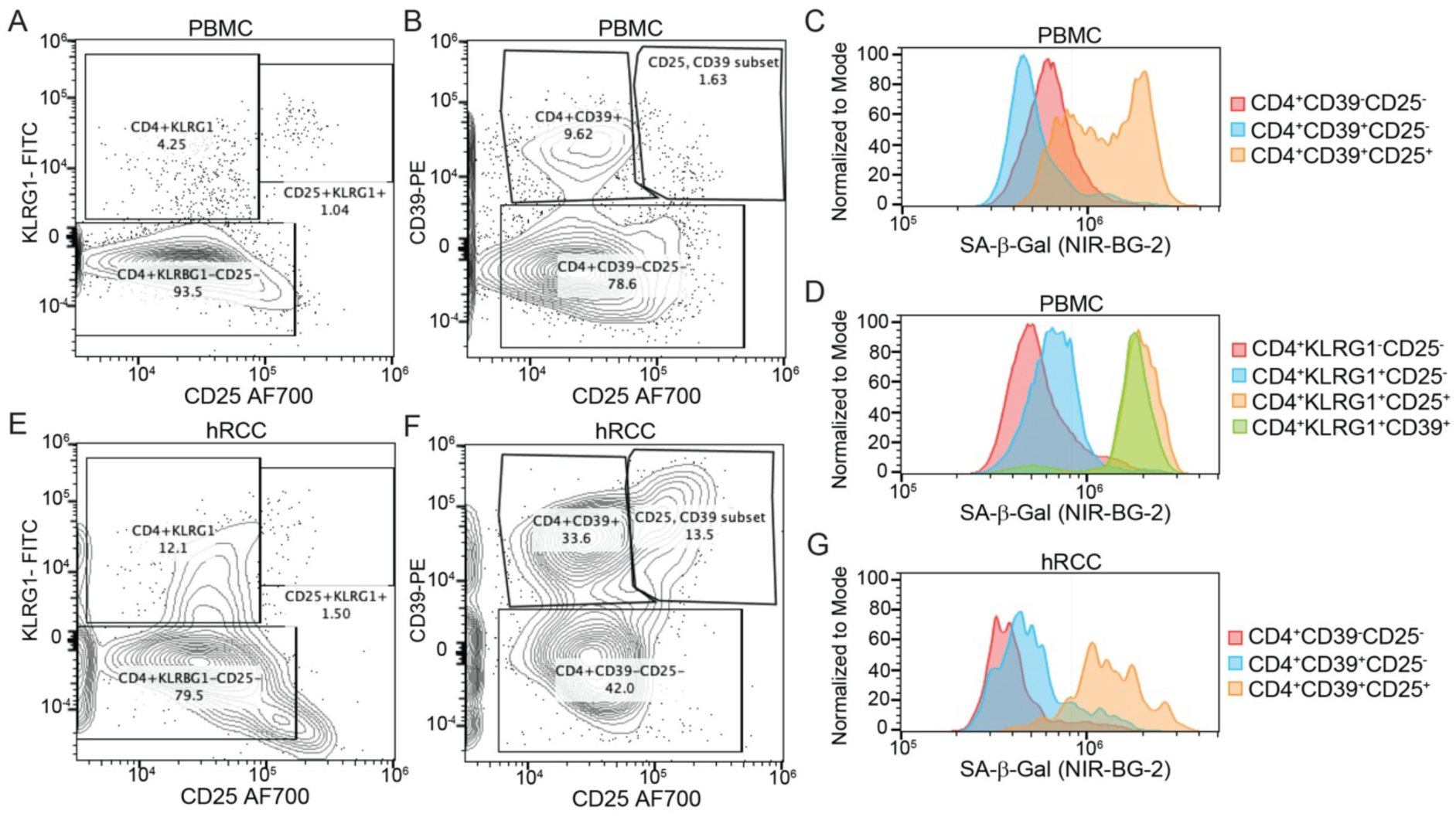
CD39 is a surrogate marker for senescent TI-Treg cells. **A.** Representative flow cytometry plots depicting KLRG1 and CD25 expression, and **B.** CD39 and CD25 expression, on CD4+ T cells from human peripheral blood mononuclear cells (PBMC). Numbers indicate the percentage of CD4+ T cells within each gate. **C.** Representative histogram showing SA-β-Gal activity, measured with the fluorogenic probe NIR-BG-2, in CD4+CD39−CD25−, CD4+CD39+CD25−, and CD4+CD39+CD25+ subsets, and **D.** in CD4+KLRG1−CD25−, CD4+KLRG1+CD25−, CD4+KLRG1+CD25+, and CD4+KLRG1+CD39+ subsets, gated as in **A.** and B. **E.** Representative flow cytometry plots depicting KLRG1 and CD25 expression, and **F.** CD39 and CD25 expression, on CD4+ tumor-infiltrating T cells from human renal cell carcinoma (hRCC). **G.** Representative histogram showing NIR-BG-2 signal in CD4+CD39−CD25−, CD4+CD39+CD25−, and CD4+CD39+CD25+ hRCC subsets, gated as in **F, C, D, G** SA-β-Gal, senescence-associated β-galactosidase; NIR-BG-2, near-infrared β-galactosidase probe. Also see supplementary Figure 5 for another replicate experiment.

In contrast, RCCs had minimal KLRG1+CD25+ Treg cells (Fig. 5e), but contained a significant number of CD39+CD25+ Treg cells (Fig. 5f), most of which were positive for NIR-BG-2 (Fig. 5g), supporting that CD39+ TI-Tregs were enriched for senescent cells. In a separate biological repeat, we found similar results as we have seen above (Fig. S5).

### Requirement of BCL-X_L_ for TI-Tregs

Our published data identified that TI-Tregs depend on BCL-X_L_ for survival ^1^. Therapeutic treatment of tumor-bearing mice with BCL-X_L_-targeting PROTACs DT2216 or PZ15227 leads to TI-Treg decrease in both human cancer specimens and mouse cancer models as well as reduction in tumor growth in various models (including RENCA RCC model, MC38 colon cancer and Py8119 breast cancer), which was attributed to significant induction of CD8+ T cell activation^1^. BCL-X_L_-targeting PROTACs, however, failed to inhibit such tumor growth in NSG mice, that lack functional T, B and NK cells.^1^ To further confirm the role of BCL-X_L_ in TI-Tregs, we crossed *Bcl2l1*-floxed mice (*Bcl2l1^fl/fl^*) in the C57BL/6J background^36^ with Foxp3^YFP/Cre^ mice (Jax: 016959)^37^ or in the Balb/C with Foxp3-Cre to generate Treg-specific *Bcl2l1*-deletion. We used RENCA RCC model for Balb/C background, the Py8119 triple negative breast cancer in the laboratory^1, 38–40^ and B16F10 melanoma. Treg-specific *Bcl2l1*-deletion led to decreased tumor growth from all three models, with RENCA model showing most significant tumor reduction (Fig. 6a) as well as survival benefit (Fig. 6b); there was a similar trend Py8119 (Fig. 6c) or B16F10 (Fig. 6d) model (Fig. 6c-6d, brown versus grey line). The Py8119 tumor reduction was also apparent in the *Bcl2l1*^fl/fl^/*Foxp3*^cre/+^ where WT and BCL-X_L_-KO Tregs co-exist at around 1:1 ratio due to the random X-chromosome imprinting (Fig. 6c, blue line).

**Figure 6.**
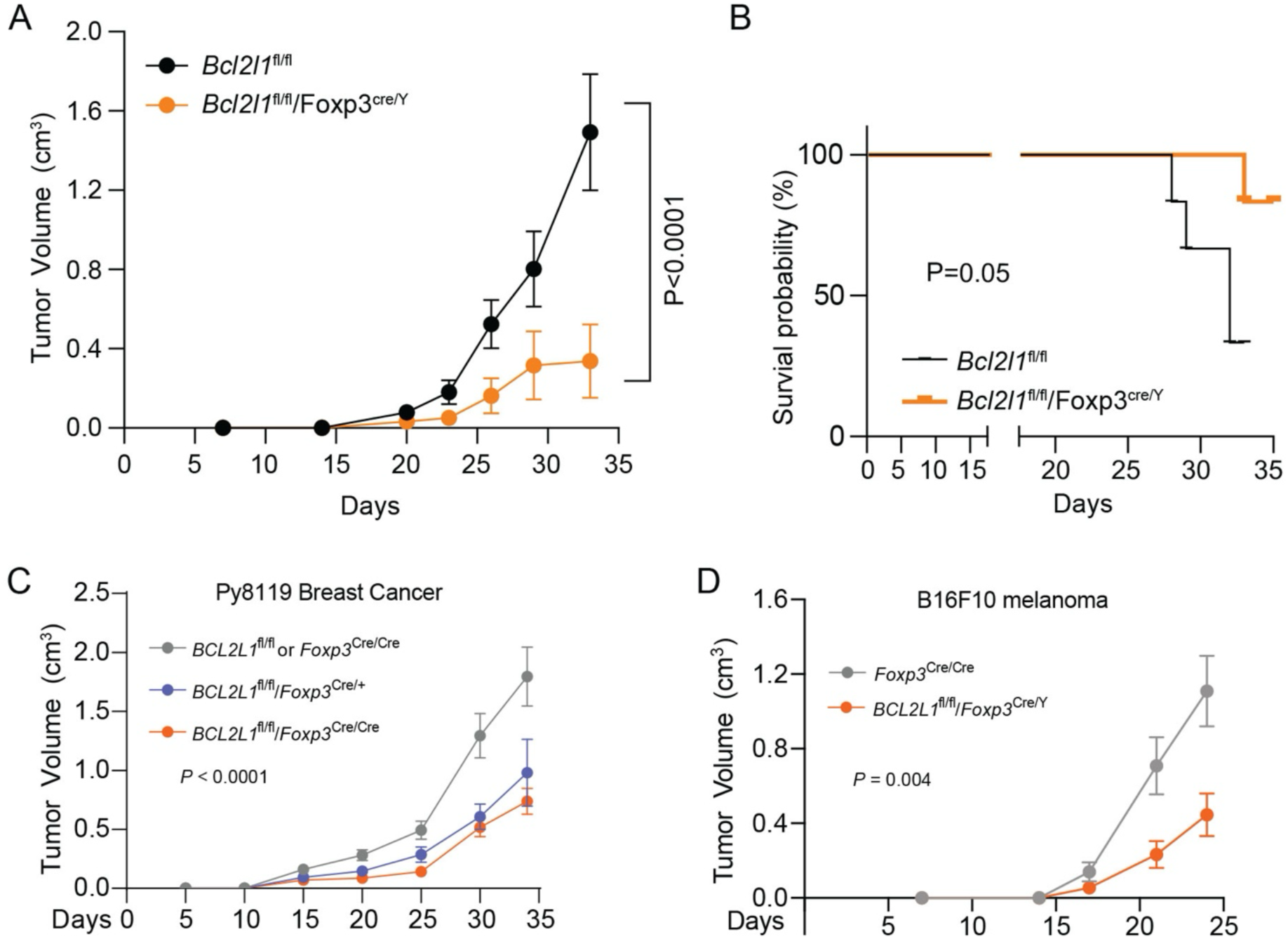
Treg-specific deletion of Bcl-XL leads to increased tumor growth in various cancer models. **A-B.** Bcl-XL deletion in Treg cells resulted in decreased Renca tumor growth. *Bcl2l1*^fl/fl^/Foxp3-cre and *Bcl2l1*^fl/fl^ 8-week old male mice received Renca cells via s.c. injection. (n=5 per group). Tumor volume (A) and overall survival (B) were shown. **C.** Bcl-XL deletion in Treg cells resulted in decreased Py8119 tumor growth. *Bcl2l1*^fl/fl^, *Bcl2l1*^fl/fl^/Foxp3^cre/cre^, or *Bcl2l1*^fl/fl^/Foxp3^cre/+^ 8-week old female mice received the engraftment of Py8119 cells. **D.** Bcl-XL deletion in Treg cells resulted in decreased B16F10 tumor growth. *Bcl2l1*^fl/fl^, *Bcl2l1*^fl/fl^/Foxp3^cre/Y^ 8-week old male mice received the engraftment of B16F10 cells. (n=5-7 per group in all tumor studies).

We confirmed the Treg-specific deletion of BCL-X_L_ in both splenic and TI Tregs within the *Bcl2l1*^ΔTreg^-tumor bearing mice, but not in other cells (Fig. 7a), supporting a critical role of BCL-X_L_ in maintaining TI-Treg survival and immunosuppressive function^1^. In agreement of immune suppressive function of BCL-X_L_^+^ Treg cells, there was a significant increase in CD8^+^ T cell activation in the tumor microenvironment of Py8119 (Fig. 7b) and RENCA (Fig. 7c), further confirming our published data using BCL-X_L_-targeting PROTACs.^1^ We identified a significant reduction of Treg cells in the draining lymph nodes from the RENCA model (Fig. 7c), supporting that Treg reduction by BCL-X_L_ depletion leads to effector T cell activation within the draining lymph nodes.

**Figure 7.**
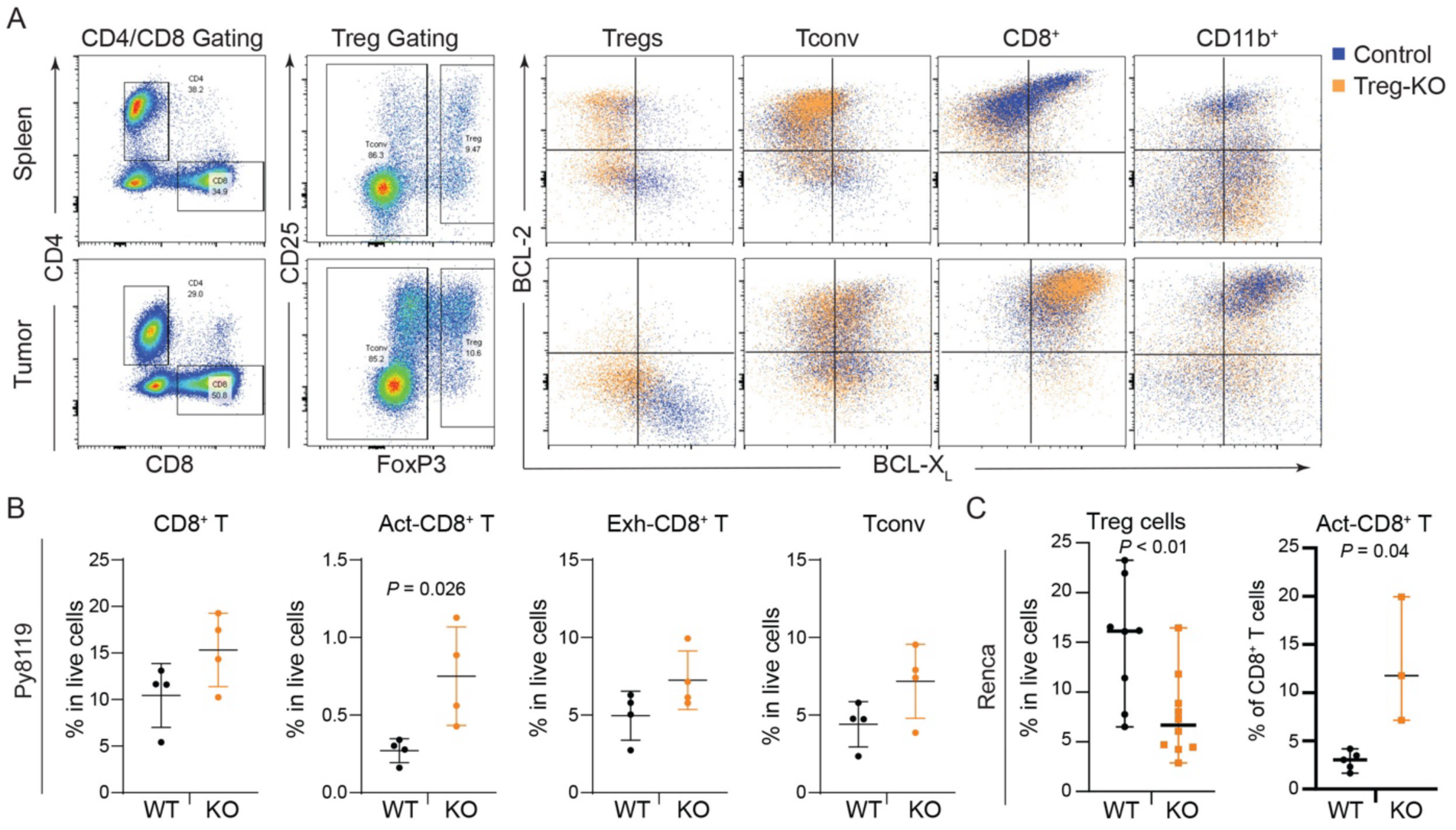
Treg-specific deletion of Bcl-XL leads to T cell activation. **A-B.** Flow cytometric analysis showing T cell gating and Treg-specific deletion of Bcl-XL in spleen and Py8119 tumors (n=4 biological repeats showing similar Bcl-XL deletion in Treg cells) (A) and significantly increased TI active GzmB+ CD8 T cells (n=4 per group) (B). **C.** Flow cytometric analysis showing that Treg-specific deletion of Bcl-XL significantly decreased Treg cells within draining lymph nodes and increased TI active GzmB+ CD8 T cells (n=3 per group) in Renca tumors.

To define the connection between CD39 and BCL-X_L_ in TI-Tregs across cancer types or in ccRCC, we performed DEGs between *ENTPD1*+ and *ENTPD1*-all cancer TI-Tregs or ccRCC TI-Tregs, as well as *BCL2L1+ or BCL2L1-*all cancer TI-Tregs or ccRCC TI-Tregs. Venn diagrams shows common DEGs across different comparisons. Please note that DEGs enriched in *ENTPD1*+ or *BCL2L1+* TI-Tregs shared core Treg signature (Fig. S6) including CCL22-Treg self-recruiting cytokine, CD177-our recently identified suppressive subtype in TI-Tregs, OX40 and GITR (TNFRSF4 and TNFRSF18, respectively)-active Treg cells, EBI3-IL35 for Treg suppression, IL1R2-core Treg gene for inhibiting inflammation etc.

### Strategic combination of senolytic depletion of TI-Treg cells and immune checkpoint inhibitors (ICI)

While the most commonly prescribed ICI is pembrolizumab, an anti-PD-1 antibody, the combination of senolytic drug DT2216 with anti-PD-1 antibody was less efficient than either drug alone in suppressing the RENCA-based tumor growth (Fig. 8a). The perplexing result is likely due to the nature of PD-1 function in T cells, which is known to — upon PD-L1 ligation — inhibit T cell receptor- and CD28-mediated signaling transduction. We treated human primary T cells ex vivo with DT-2216, Anti-PD-1 antibody, or combination, and found that the combination significantly increased annexin V+ apoptotic CD8^+^ T cells (Fig. 8b). Instead, we combined BCL-X_L_-targeting PROTAC PZ15227 with anti-PD-L1 antibody to treat mice bearing RENCA tumors, with the reasoning that anti-PD-L1 antibody doesn’t directly bind to T cells and won’t sensitize T cells to BCL-X_L_-depletion. We did find that the combination of PZ15227 with anti-PD-L1 antibody inhibited RENCA tumor growth more efficiently than either drug alone (Fig. 8c). Profiling the tumor infiltrating lymphocytes identified the differential effect of PZ15227 and anti-PD-L1 on TI-CD8+ T cells: 1) PZ15227 treatment leading to more active Gzmb^+^ CD8^+^ T cells in both spleen and tumors; 2) anti-PD-L1 treatment resulting in increased number of TI-CD8+ T cells in tumors; and 3) combination treatment showing additive effect in CD8+ T cells in tumors while maintaining the ratio of active Gzmb^+^ CD8^+^ T cells in both spleen and tumor (Fig. 8d-8e). This data supports the combinatory regimen of using senolytic BCL-X_L_-targeting PROTAC and anti-PD-L1 antibody.

**Figure 8.**
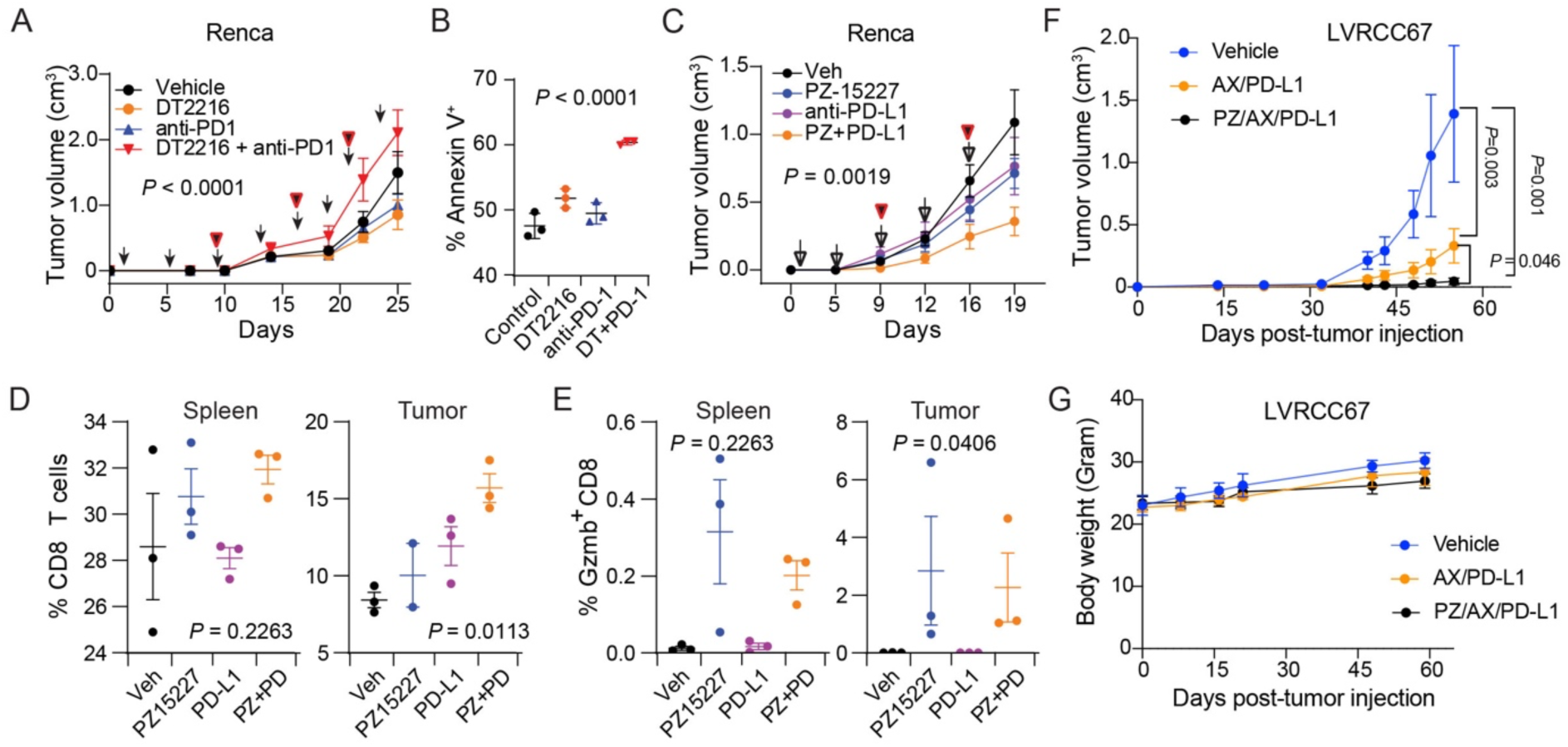
Rational design therapeutic modality for combinatory therapy using Bcl-XL-targeting PROTAC and ICIs. **A.** Bcl-XL-targeting PROTAC and anti-PD-1 antibody should not be combined for therapy. Balb/C mice bearing Renca tumors were treated with vehicle, DT2216, anti-PD1 antibody, or co-treatment of DT2216 and anti-PD1 antibody (n=5 per group). **B.** DT2216 and anti-PD1 antibody combination induced apoptosis of T cells. Splenic CD8+ T cells were activated with anti-CD3/anti-CD28 dynabeads for two days, following with treatment of control, 50nM of DT2216, 100 ng/ml anti-PD-1, or combination (n=3 biological replicates). Annexin V staining was used to differentiate apoptotic cells. **C-E**. Bcl-XL-targeting PROTAC and anti-PD-L1 antibody exhibit complementary effect in T cell activation and tumor suppression. Balb/C mice bearing Renca tumors were treated with vehicle, PZ15227, anti-PD-L1 antibody, or co-treatment of PZ15227 and anti-PD-L1 antibody (n=5 per group). Tumor volume was recorded and graphed **(A)** (n=5 per group); Splenic and TI-CD8 T cells were enumerated **(B)** with GzmB+ to depict active CD8 T cells **(C)** (n=3 per group)**. F-G.** Combination of Bcl-XL-targeting PROTAC and standard frontline ccRCC therapy led to near complete tumor eradication. C57BL/6J male mice carrying LVRCC67 tumors were treated with vehicle, axitinib+anti-PD-L1, or PZ15227+ axitinib+anti-PD-L1. Tumor volume was recorded and graphed **(F)** as well as body weight **(G)** (n=5 per group). Also see supplementary Figure 6 showing single Bcl-XL-targeting PROTAC treatment did not inhibit LVRCC67 tumor growth in vivo.

In a more pathological relevant ccRCC model, we used LVRCC67 that was generated from a primary ccRCC tumor carrying *Vhl*, *Tp53* and *Rb1 deletions*. PZ15227 alone didn’t inhibit tumor growth in the LVRCC67 model, neither another BCL-X_L_-targeting PROTAC WH244^41^ with known in vivo activity (Fig. S7). We combined PZ15227 with standard front line therapy axitinib + anti-PD-1 therapy in the LVRCC67 model with slight modification by replacing anti-PD-1 with anti-PD-L1 based on above data. Axitinib+anti-PD-L1 antibody significantly inhibited tumor growth of LVRCC67 in the C57BL/6J mice; addition of PZ15227 nearly completely eradiated tumor growth (Fig. 8f). All three groups exhibited similar body weight gain, validating the safety profile of the triple treatment in vivo (Fig. 8g).

## Discussion

Apoptosis of Treg cells have been demonstrated as a major mechanism of immune suppression within the TME of several cancer types, such as ovarian^34^ and hepatocellular carcinomas^42^, accompanied with CD177 positivity as we have shown in breast and renal cancers.^14^ It is believed that the oxidative stress within the TME is the cause of Treg cell apoptosis, which leads to the release of intracellular ATP via the adenosine A_2A_ receptor and subsequent catabolism of ATP to adenosine - an immune suppressive metabolite via upregulation of CD39 in apoptotic TI-Tregs.^34^ However, apoptotic cells under physiological conditions are removed instantaneously and hence these cells need to be constantly replaced – a relatively less efficient way of maintaining the immune suppressive TME.

Apoptosis and senescence are two overlapping yet distinct pathways that are essential to eliminate cancer cells and halt cancer cell proliferation, respectively during many chemo- or radiotherapeutic treatment. Apoptotic signature and senescence signature are loosely correlative in different clusters of Treg cells (Fig. S3b). Normally apoptotic induction is due to severe DNA damage that cannot be repaired; whereas low/intermediate level of DNA damage results in elevated and sustained cell cycle inhibitor due to p53-p21 (CDKN1A) axis and the consequent permanent cell cycle arrest – senescence. The distinct gene signature of senescence includes Glb1 expression and SA-β-GAL activation, SASP phenotype, as well as sustained co-elevation both pro- and anti-apoptotic BCL2 family members. Hence we believe senescence-like Treg cells are the product of moderate oxidative stress within the TME as we have shown (**Fig. S4b**). Interestingly, both apoptotic Treg cells – as reported and senescence-like Treg cells – as we showed here converge on the same immune suppressive pathways, *i.e.* CD39-mediated ATP to adenosine conversion. Interestingly, aging-associated Treg cell senescence results in the loss of immune suppressive function, a phenotype that agrees with the increased inflammation under physiological aging.

Envisioning that apoptosis and senescence are two sides of the same coin for TI-Treg cells, it is perceivable that senescent and apoptotic Tregs work together in the TME to achieve stronger immune suppression. While purification of apoptotic Treg cells from solid tumors are nearly impossible, most conclusions have to be drawn from in vitro induced apoptotic cells either by oxidative stress, anti-Fas antibody or irradiation. Purification of senescence-like Treg cells faces the same technical difficulty as there is no single marker that can define senescence. KLRG1 has been established as a marker for T cell senescence, which is validated in our data as well for labeling Treg cell senescence in PBMC (**Fig. 5**); however, KLRG1 fails to faithfully label senescence-like Treg cells in the human RCCs. CD39 is a much more specific marker for labeling senescence-like TI-Treg, with more than 90% CD39^+^ TI-Treg cells have active SA-β-GAL (**Fig. 5**). CD39^+^ conventional CD4+ T cells, however, exhibit only minimal positivity for NIR-BG-2 staining. Interestingly, CD39 expression in TI-Treg cells seem to be uniform phenotype across mouse and human tumors, which can be induced from PBMC Treg cells by RCC-coculture by H2O2 induced oxidative stress. CD39+ TI-Treg cells express high levels of senescence signature genes, immune suppressive and active Treg genes, as well as elevated dependence on BCL-X_L_, providing a unique senolytic depletion of senescence-like Treg cells within the TME as we have shown before. In addition, targeting BCL-X_L_ with PROTAC may also push TI-Treg cells – especially those clusters with enriched apoptotic module (Fig. S3b) to their inevitable death and eventually lose their suppressive function.

Interestingly, TI-Treg senescence-like state provides an interesting angle for guiding the design of therapeutic regimen using senolytic drugs and ICIs. Our data argue against random selection of ICI drugs for combination with BCL-X_L_-degrading PROTACs due to potential additive effects on CD8+ T cell toxicity. Combination of BCL-X_L_-degrading PROTAC and anti-PD-1 antibody leads to significant apoptosis of CD8+ T cells and cancelling each other’s inhibitory effect in tumor growth. Anti-PD-L1, however, works additively with BCL-X_L_-degrading PROTAC to inhibit tumor growth. One potential explanation of the differential effect lies in the different mechanisms of action, with anti-PD-1 antibody directly binding to PD-1 on T cells but anti-PD-L1 antibody binds to PD-L1 on tumor cells or myeloid cells in the TME. The direct ligation of anti-PD-1 antibody to PD-1 sensitizes T cells to BCL-X_L_-degrading PROTACs. Interestingly, the senolytic depletion of BCL-X_L_-dependent TI-Treg cells complements with anti-PD-L1 antibody treatment; the former results in T cell activation while the latter is more involved in an increase of T cell number in the TME.

## Materials and Methods

### Chemical synthesis and quality control

DT2216 and PZ15227 were synthesized in-house following previously published procedures,^43, 44^ whereas WH244 was prepared according to the synthetic procedure disclosed in patent WO2023107606A1. The identity of the synthesized compounds was confirmed by appropriate analytical methods, including MS and NMR spectroscopy. All compounds used for biological studies had a purity of ≥95% as determined by analytical HPLC.

For in vitro studies, compounds were prepared as stock solutions in DMSO and diluted into appropriate culture medium before use. For in vivo studies, compounds were formulated in 50% Phosal 50 PG / 45% Miglyol 810N / 5% Polysorbate 80 and administered as described in the corresponding animal experiments.

### Mouse study

All animal experiments were conducted under protocols approved by the University of Florida Institutional Animal Care and Use Committee (IACUC201810399; IACUC202400000275; and IACUC202200000379). Mice were maintained under specific pathogen-free (SPF) conditions in an Association for Assessment and Accreditation of Laboratory Animal Care (AAALAC)-accredited animal facility at the University of Florida, with experimental animals housed in the same room. All procedures were carried out in compliance with institutional IACUC requirements. At experimental endpoints, animals were euthanized by inhalation of medical-grade carbon dioxide in accordance with current American Veterinary Medical Association (AVMA) recommendations and University of Florida IACUC policies. Temperature is set at 22.2 °C with 21.1–23.3 °C daily fluctuation of 2 °C. The humidity set point is 50% with ±15% daily variation. 12:12 hrs photoperiod was maintained with light intensity range of 15.6–25.8 FC.

*Bcl2l1*^fl/fl^ mice in the C57BL/6J genetic background was kindly provided from Hennighausen,^36^ which was crossed with Foxp3^cre^ in C56BL/6J genetic background (JAX #016959)^37^ to produce *Bcl2l1*^fl/fl^, *Bcl2l1*^fl/fl^*/Foxp3^cre/cre^*, or *Bcl2l1*^fl/fl^*/Foxp3^cre/+^*female mice for engraftment of 100,000 Py8119 cells; or *Bcl2l1*^fl/fl^ and *Bcl2l1*^fl/fl^*/Foxp3^cre/Y^* male mice for the engraftment of 200,000 of B16F10 cells. *Bcl2l1*^fl/fl^ mice were also backcrossed to Balb/C genetic background for 8 generations and further crossed with a different Foxp3-cre strain of Balb/C generated by Dr. Sakaguchi for two more generations.^45^ *Bcl2l1*^fl/fl^/Foxp3-cre and *Bcl2l1*^fl/fl^ male mice were used for receiving 100,000 RENCA cells. Tumors and body weight were monitored twice weekly and recorded according.

### Human tissue collection and cell isolation

Fresh human clear cell renal cell carcinoma (ccRCC) specimens and matched peripheral blood samples were obtained from patients undergoing surgical resection following informed consent. Samples were provided in de-identified form through the University of Florida Biorepository and Tissue Procurement Core. De-identified peripheral blood samples from healthy donors were obtained from LifeSouth Community Blood Centers. Peripheral blood mononuclear cells (PBMCs) were isolated by density-gradient centrifugation using Ficoll-Paque (GE Healthcare), with SepMate tubes (STEMCELL Technologies). All usages of human specimens were approved by the University of Florida Institutional Review Board (IRB) under IRB number: IRB201901677.

Fresh tumor specimens were transferred to the research laboratory following surgery and processed by enzymatic digestion with 1 mg/mL collagenase type VI and 50 μg/mL DNase I, combined with mechanical dissociation using a gentleMACS dissociator (Miltenyi Biotec). Cell suspensions were passed through a 100μm cell strainer, and red blood cells were lysed using ammonium chloride potassium (ACK) lysis buffer. Treg cells were magnetically enriched using a human CD4⁺CD127^lowCD25⁺ regulatory T-cell isolation kit (STEMCELL Technologies).

### Direct coculture of regulatory T cells with renal cancer cells

Treg cells were directly cocultured with 769-P renal cell carcinoma cells or HK-2 renal proximal tubular epithelial cells in six-well plates at a Treg-to-RCC cell ratio of 2:1. Cocultures were maintained in RPMI-1640 supplemented with 10% fetal bovine serum, 100 μg/mL streptomycin, 100 U/mL penicillin, 1 mM sodium pyruvate, 10 mM HEPES, and 30 IU/mL recombinant human interleukin-2 (IL-2). Cells were incubated at 37 °C in a humidified atmosphere containing 5% CO₂ for 72 hrs, when Treg cells were recovered for downstream analyses.

### Treg suppression assay

Naïve CD4⁺ T cells were isolated from human PBMCs using the Naïve CD4⁺ T Cell Isolation Kit II (Miltenyi Biotec; catalog no. 130-094-131). Cells were stimulated overnight with anti-CD3/CD28 Dynabeads (Thermo Fisher Scientific) in the presence of 30 IU/mL recombinant human IL-2 (R&D Systems).

Pre-activated responder CD4⁺ T cells were labeled with 5 μM CellTrace Violet in 1 mL phosphate-buffered saline (PBS) for 15 min and subsequently cocultured with Treg cells recovered from 769-P or HK-2 cocultures. Suppression assays were performed in 96-well round-bottom plates at responder-to-Treg ratios of 2:1, 4:1, and 8:1 in a final volume of 200 μL per well. Cultures were maintained for 72 h with continued anti-CD3/CD28 Dynabead stimulation at a bead-to-cell ratio of 1:1. Responder-cell proliferation was assessed by flow cytometric analysis of CellTrace Violet dilution.

### Flow cytometry

Mouse tumors were digested into single cell suspensions, following with surface staining and intracellular staining for immune profiling different immune cell compartments.^1, 14, 46^

Single-cell suspensions prepared from human ccRCC specimens were washed and incubated in complete RPMI-1640 containing 100 μM bafilomycin A1 for 1 h at 37 °C, followed by incubation with 5 μM NIR-BG2 for an additional 1 h at 37 °C. Cells were washed twice with PBS and stained with Zombie Aqua Fixable Viability Dye (BioLegend) at a dilution of 1:500.

Surface immunophenotyping was performed using FITC-conjugated anti-human KLRG1 (clone SA231A2), BV605-conjugated anti-human CD4 (clone RPA-T4), Spark NIR 685-conjugated anti-human CD45 (clone 2D1), PE/Cyanine7-conjugated anti-human CD25 (clone M-A251), BV421-conjugated anti-human CD39 (clone A1), and APC-conjugated anti-human CD127 (clone A019D5). All antibodies were used at a dilution of 1:100. A human Fc receptor-blocking reagent was used to minimize nonspecific binding. Samples were acquired using an Aurora spectral flow cytometer (Cytek Biosciences), and data were analyzed using FlowJo software.

### Bulk RNA sequencing and bioinformatic analysis

Total RNA was extracted from Treg cells recovered from 769-P or HK-2 cocultures and submitted to the University of Florida Interdisciplinary Center for Biotechnology Research Gene Expression and Genotyping Core (RRID: SCR_019145). RNA concentration was measured using a Qubit fluorometer, and RNA integrity was assessed using a Bioanalyzer RNA assay. Samples with an RNA integrity number of at least 8 were used for library preparation.

Polyadenylated RNA was enriched using the NEBNext Poly(A) mRNA Magnetic Isolation Module, and sequencing libraries were prepared using a NEBNext Ultra II library preparation kit (New England Biolabs). Libraries were sequenced on an Illumina NovaSeq 6000 platform to generate 150-bp paired-end reads.

Transcript abundances were quantified using kallisto by pseudoalignment to a human reference transcriptome corresponding to GRCh38. Differential expression between Treg cells recovered from 769-P and HK-2 cocultures was assessed using the limma package in R.^47^ Gene set enrichment analysis^48^ was performed to assess differences in biological pathways between the two conditions. Figures were generated using ggplot2 in R.

### Pan-cancer tumor-infiltrating lymphocyte atlas

Treg analysis was performed on the v1.0.1 uTILity atlas, a harmonized collection of publicly available tumor-infiltrating lymphocyte scRNA-seq with paired TCR sequencing. The release contains 2,606,012 cells from 682 samples, with 251 patients, across 127 studies and 13 cancer types. Cells were removed if they carried over 10% mitochondrial reads, over 60% ribosomal reads, fewer than 100 detected features, or a feature count more than 2.5 standard deviations from the run mean. Doublets were called per run with scDblFinder (v1.22.0)^49^ and only singlets retained. The processed output of uTILity was loaded into R as a Seurat v5.5.0 object^50^. Tregs were isolated using the consensus annotation from the Azimuth v0.5.0^51^ and SingleR v1.22.0^52^ pipelines, with the Monaco^53^ and HPCA^54^ reference data sets and expression of FOXP3 or IL2RA. The resulting cells were further filtered for expression of CD3D, CD3E, TRAC, or CD2 to ensure retention of only T cells.

Atlas Tregs were re-embedded on 2,000 variable genes and 30 principal components with Harmony v2.0.5^55^, using the sample of origins as a variable. TCR gene segments, immunoglobulin segments, ribosomal and mitochondrial genes were excluded from the variable features. Downstream dimensional reduction and clustering utilized the 30 Harmony-projected PCA dimensions. Louvain clustering was based on stability, using 15 bootstraps of 80% of the cells, resulting in a resolution of 0.4. UMAP embedding utilized the uwot (v0.2.4) defaults.

### ccRCC Single-Cell Analysis

The ccRCC arm used GSE121638^15^. In Seurat, cells were filtered for 200 to 5000 detected genes with < 10% mitochondrial reads, integrated across patients by CCA, and clustered as above using the 20 correct CCA dimensions with the Louvain approach at a resolution of 1.2. Tregs were identified using the previously described consensus annotation and compartment-specific FOXP3 thresholding, in which a cluster was defined as a Treg when its FOXP3 detection rate exceeded three times the background rate of all other T cell clusters in the same compartment, subject to an absolute floor of 10% and to IL2RA detection above the object-wide median.

### Senescence scoring

Senescence signatures were SenMayo^56^, an aging-Treg signature derived from Guo et al.^29^ by mouse-to-human ortholog mapping, and the Reactome, GO:BP and Fridman senescence sets^57^. Senescence was additionally decomposed into six independent axes: cell cycle arrest, T cell senescence, and the SASP chemokine, cytokine, protease and growth factor modules. Pathway collections for enrichment testing were MSigDB Hallmark, GO:BP and Reactome, restricted to sets of 10 to 500 genes. Signatures were scored per cell with UCell algorithm^58^ in escape v2.6.2^15^, a rank-based method that is robust to the sparsity and variable depth of droplet data. Scores were then revisualized on log detected genes.

### Differential expression

Differential expression analysis used pseudo bulk with patients as the unit of replication. Counts were summed within patients and groups. Patients contributing at least 10 cells to both groups were retained. Genes were filtered using edgeR (v4.6.3)^59^. Libraries were normalized by trimmed mean of M-values, and the contrast was fitted with voom in limma v3.64.3^60^ using empirical Bayes moderation on the design ‘∼ patient + group’. Gene set enrichment^48^ used fgsea v1.34.2 on genes ranked by signed -log10(p) rather than by fold change. Enrichment was computed against Hallmark, GO:BP, and Reactome, restricted to sets of 10 to 500 genes and to genes detected in the data. Significance was Benjamini-Hochberg adjusted at 0.05.

### Trajectory

Pseudotime was fitted with slingshot (v2.16.0)^61^ on the Harmony embedding, capped at 40,000 cells. The lineage graph was fitted on a coarser clustering (resolution 0.2) than the rest of the analysis. The root was chosen by combining the peripheral blood fraction with a naive Treg marker score (CCR7, SELL, TCF7, LEF1, KLF2, IL7R, S1PR1), and both components are reported. Signature trends along pseudotime were fitted with generalized additive models in mgcv v1.9-4 and the reported direction compares the last quintile against the first so that a non-monotone fit is not described as rising throughout.

### TCR repertoire

Repertoires were assembled with scRepertoire v2.6.2^62^ using strict clonotype definitions over paired TRB and TRA. Clonal expansion by cluster was tested against the rest of the Tregs, and clonal sharing along the trajectory was tested by permutation with 2,000 iterations.

## Data Availability and Code Availability

All code for the single-cell analysis is available at https://github.com/BorchLab/SenescentTregs. The data is available at https://zenodo.org/records/10211240.

## Author Contributions

Study conceptualization and design: T.O., N.B., and W.Z. Experimentation, materials, data acquisition and analysis: T.O., N.B., M.K., C.K.M., W.C.T., K.W. Chemical synthesis and formulation: P.Z. and G.Z. Supervision and scientific interpretation of data: M.K., R.K. and W.Z. Writing – original draft: W.Z. and T.O. Writing – review & editing: All authors. Funding Acquisition: W.Z. All authors have read and agreed to the published version of the manuscript.

## Acknowledgements

The work is partly supported by NIH/NCI: CA269661 (W. Zhang), CA260239 (W. Zhang; D. Zhou; G. Zheng), and CA290792 (W. Zhang; G. Zheng; K.S.M. Smalley), University of Florida Health Cancer Institute grant P30CA247796 (PI: T. George), an endowment fund from the Dr. and Mrs. James Robert Spenser Family (W. Zhang).

---

**Supplementary Figure 1.**
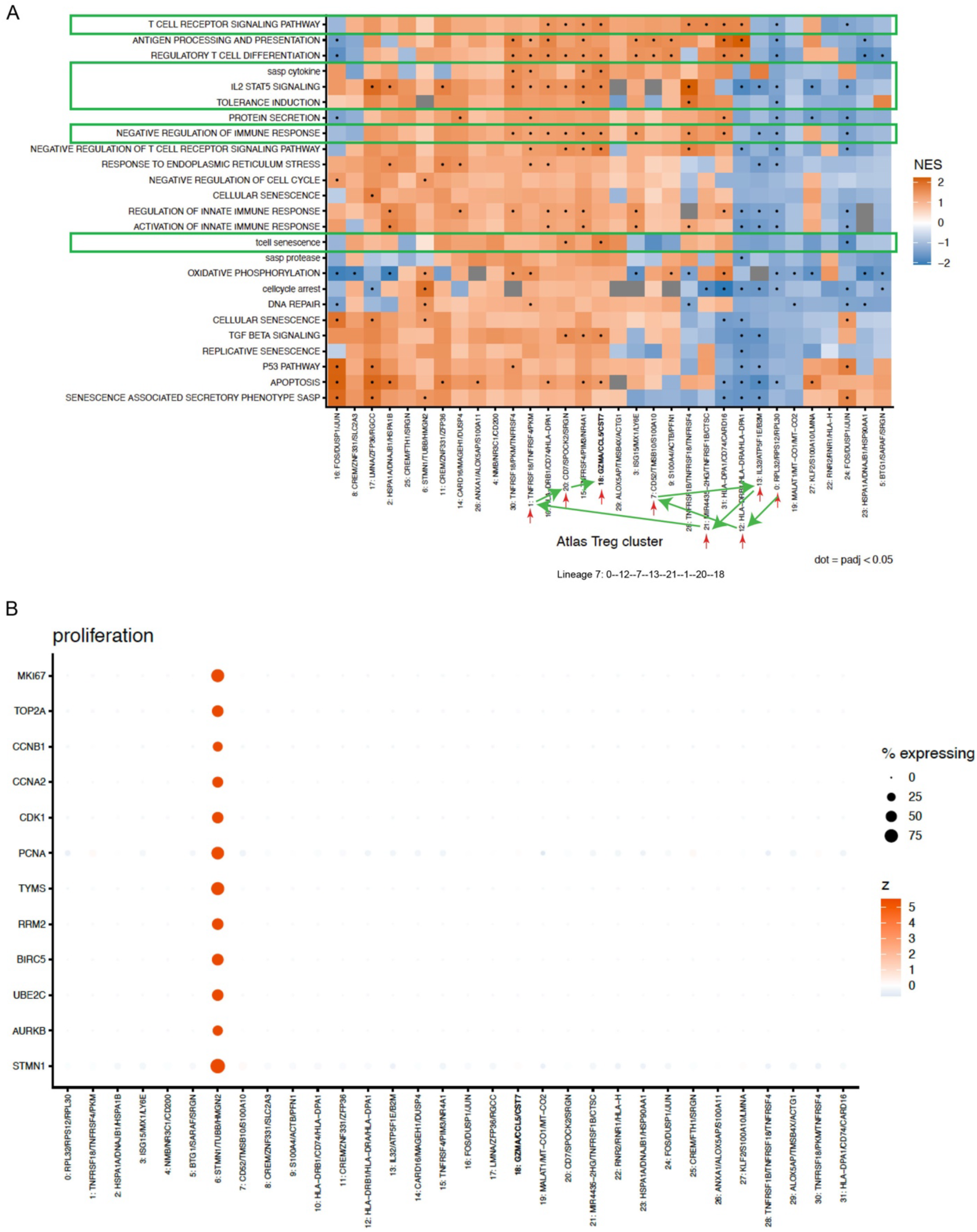
Human Cancer TI-Treg cells exhibit various immune suppressive states. **A.** Heatmap showing GSEA analysis of all Treg clusters with green arrows showing the lineage 7 trajectory. **B.** Dotplot showing gene expression patterns related to proliferative signatures across all Treg cell clusters.

**Supplementary Figure 2.**
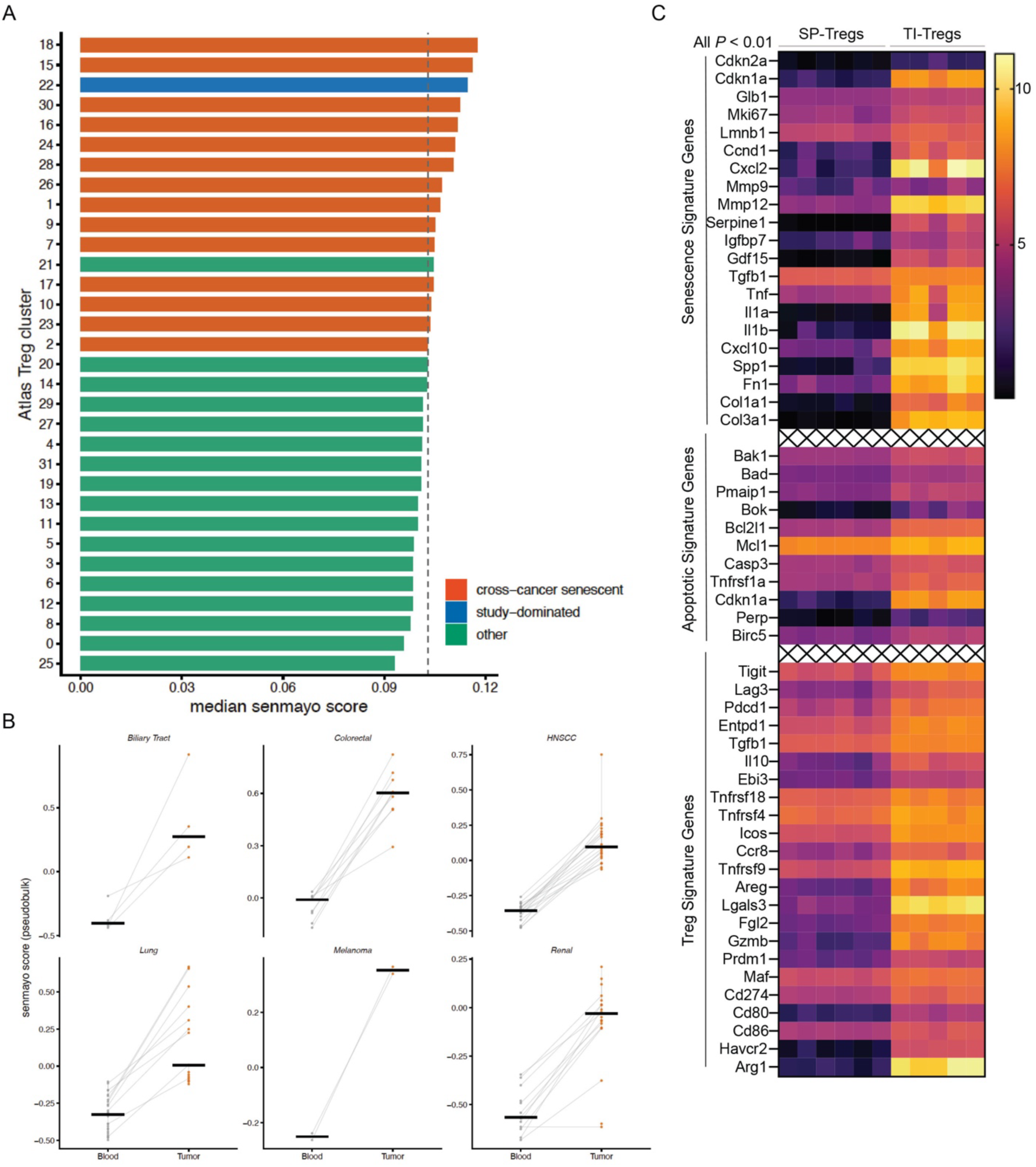
Senescence-like state of TI-Treg cells exhibits distinct set of immune suppressive signature. **A.** Bar graph showing median senmayo senescence score of all Treg clusters. **B.** Various cancers exhibit increased senmayo senescence score relative to their blood Treg counterparts, showing biliary track cancers, colorectal cancers, HNSCC, lung cancers, melanoma, and renal cancers. **C.** Enrichment of senescence related genes with TI-Tregs relative to splenic Tregs from MC38-tumor bearing mice (n=5-6 biological replicates).

**Supplementary Figure 3.**
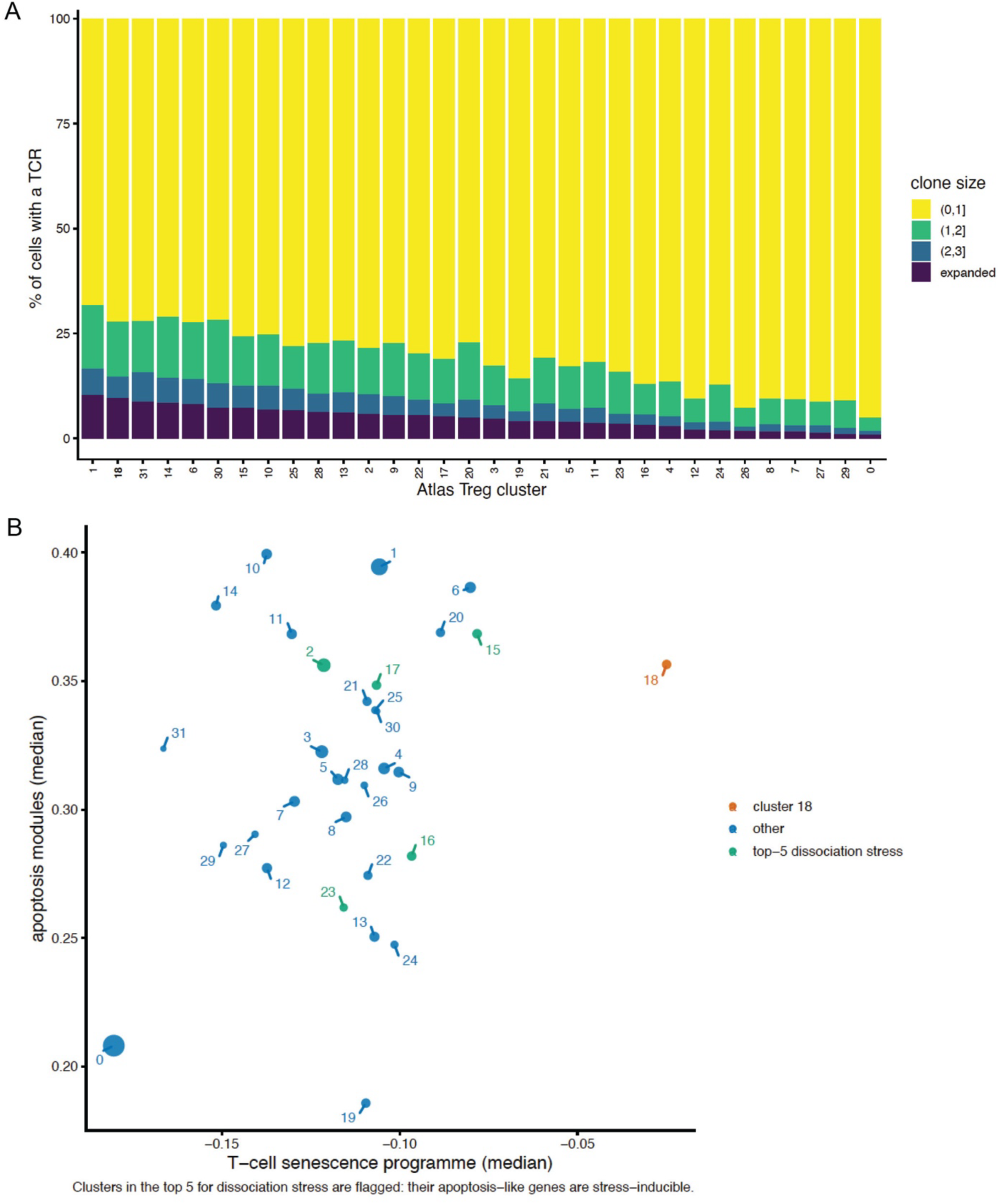
TI-Treg cells exhibit various level of clonal expansion, apoptosis, as well as senescence signatures across all clusters. **A.** Clonal expansion of Treg cells with all different clusters, noting that cluster 18 ranked 2^nd^ in clonally expanded Treg cells. **B.** Correlation of apoptotic module and T-cell senescence module across all Treg cell clusters.

**Supplementary Figure 4.**
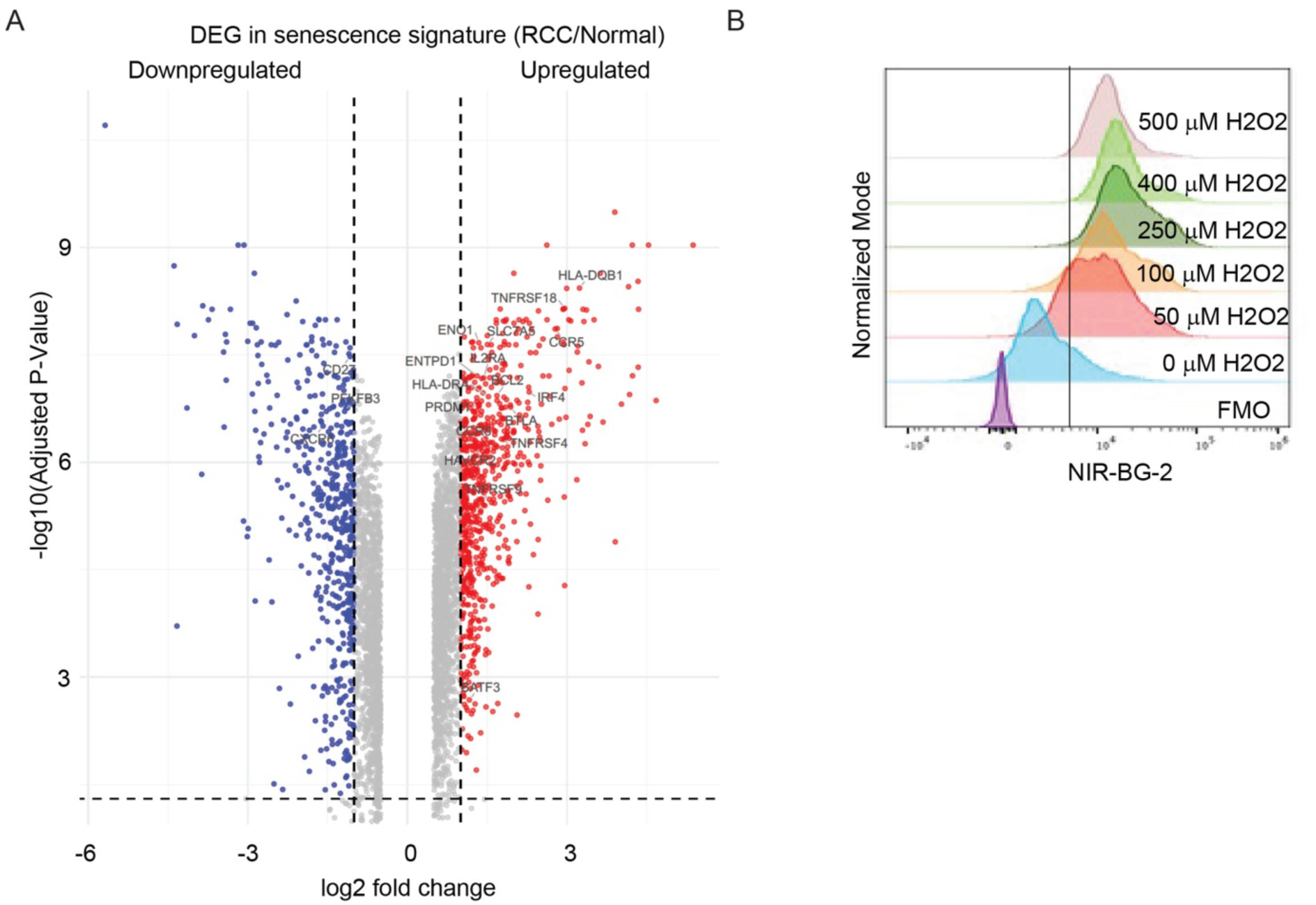
Renal cancer cells induce Treg senescence and enhance Treg suppressive function. **A.** Volcano plot highlighting Treg activation and suppression-associated genes upregulated in Tregs following with co-culturing with 769-P cells compared with HK-2 cells for 72 h. Dashed lines indicate log₂ fold-change thresholds of ±1 and an adjusted **P** value threshold of 0.05. Upregulated and downregulated genes meeting these thresholds are shown in red and blue, respectively; other genes are shown in grey. Selected genes of interest are labeled. **B.** Flow cytometry histograms showing NIR-BG-2 fluorescence following treatment with the indicated concentrations of hydrogen peroxide (H₂O₂). Histograms are normalized to mode. FMO, fluorescence-minus-one control.

**Supplementary Figure 5.**
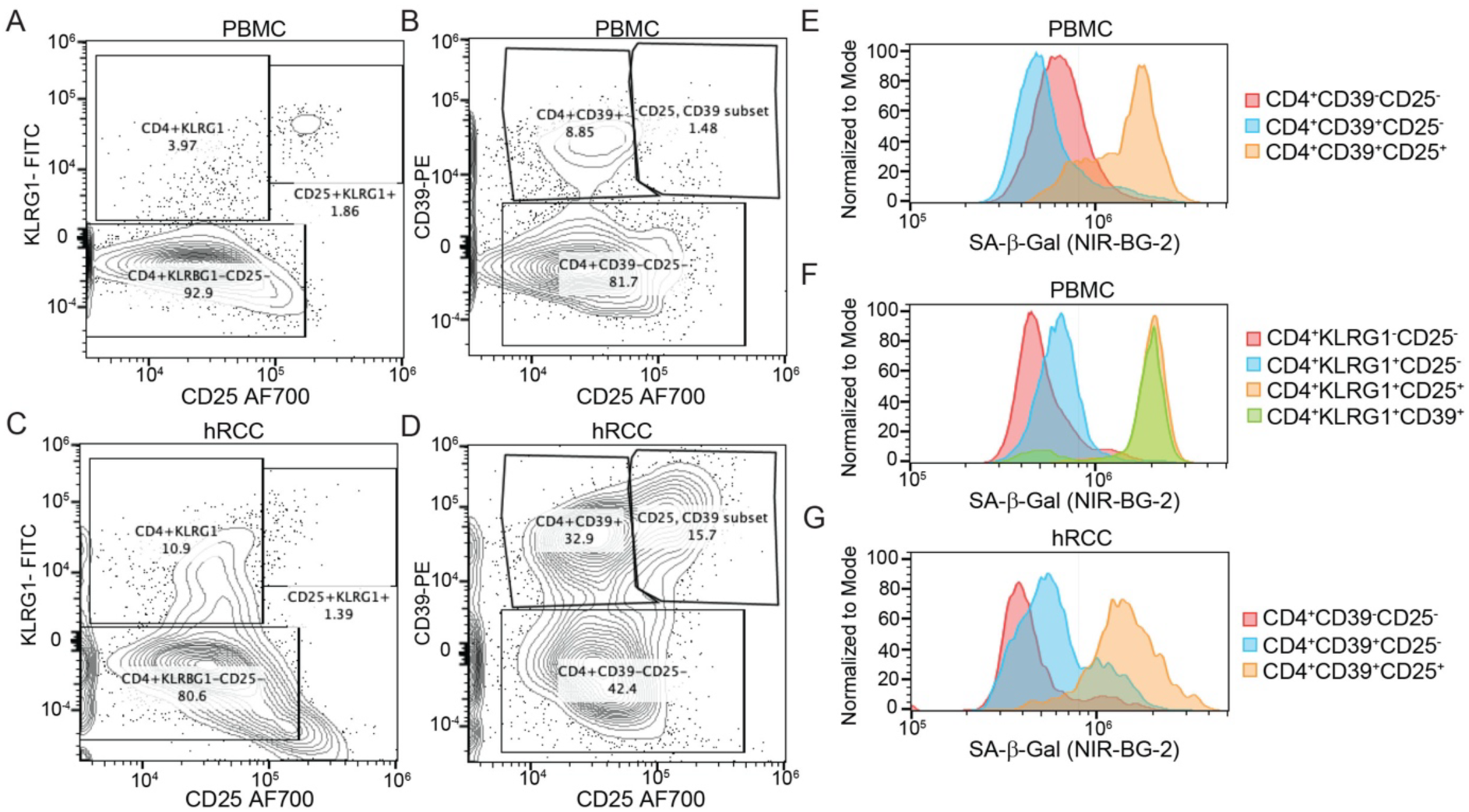
CD39 is a surrogate marker for senescent TI-Treg cells. **A.** Representative flow cytometry plots depicting KLRG1 and CD25 expression, and **B.** CD39 and CD25 expression, on CD4+ T cells from human peripheral blood mononuclear cells (PBMC). Numbers indicate the percentage of CD4+ T cells within each gate. **C.** Representative histogram showing SA-β-Gal activity, measured with the fluorogenic probe NIR-BG-2, in CD4+CD39−CD25−, CD4+CD39+CD25−, and CD4+CD39+CD25+ subsets, and **D.** in CD4+KLRG1−CD25−, CD4+KLRG1+CD25−, CD4+KLRG1+CD25+, and CD4+KLRG1+CD39+ subsets, gated as in **A.** and B. **E.** Representative flow cytometry plots depicting KLRG1 and CD25 expression, and **F.** CD39 and CD25 expression, on CD4+ tumor-infiltrating T cells from human renal cell carcinoma (hRCC). **G.** Representative histogram showing NIR-BG-2 signal in CD4+CD39−CD25−, CD4+CD39+CD25−, and CD4+CD39+CD25+ hRCC subsets.

**Supplementary Figure 6.**
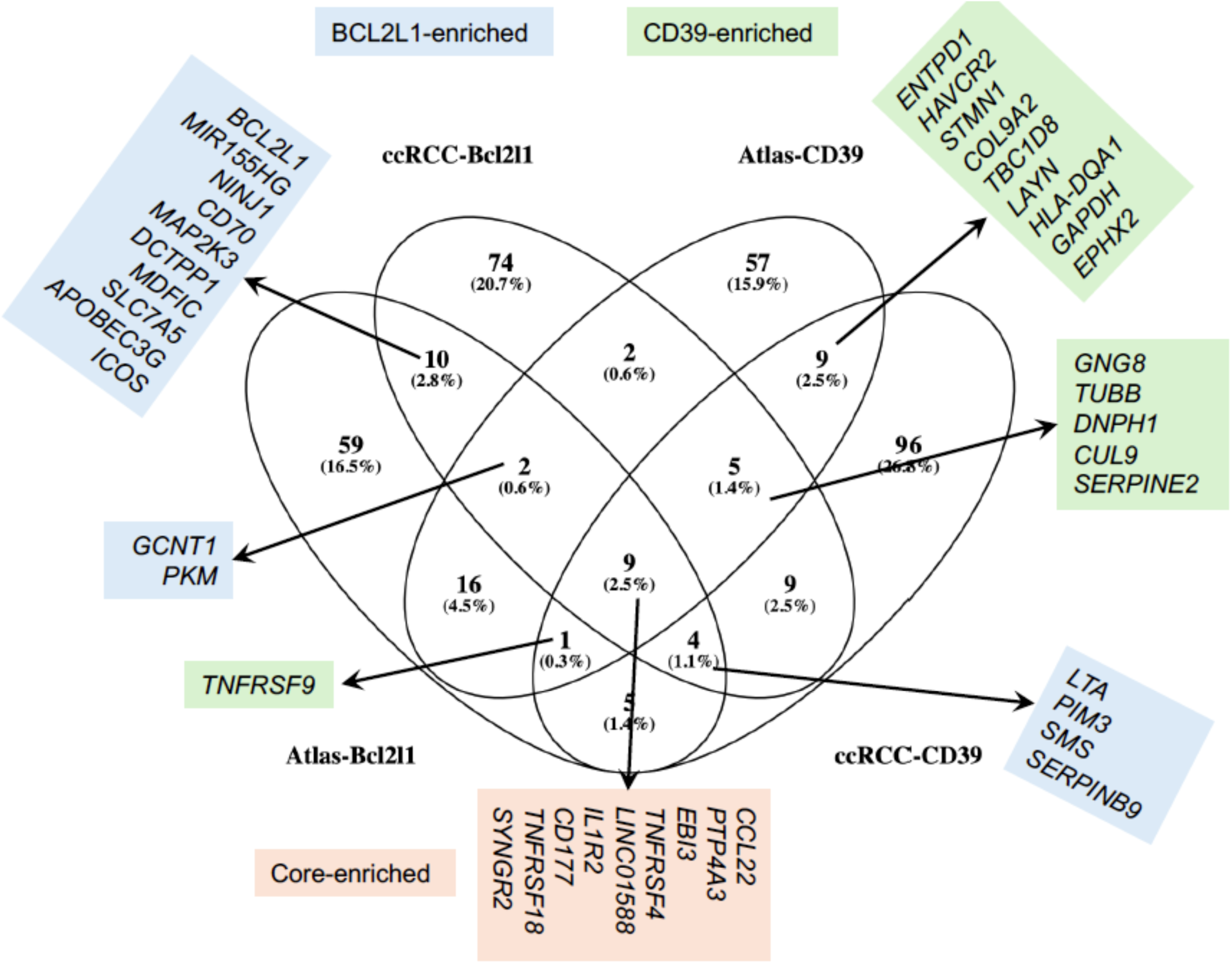
Enrichment of core Treg related gene signatures among *ENTPD1 (CD39)*+ and *BCL2L1*+ TI-Treg cells. Pan-cancer TI-Treg cells and ccRCC TI-Tregs were separated by *ENTPD1+ and ENTPD1-, or by BCL2L1*+ and *BCL2L1*-, with DEGs analyzed. Venn diagram showing common DEGs elevated in *ENTPD1+* or in *BCL2L1*+ TI-Tregs. Please note a core of TI-Treg signature is enriched.

**Supplementary Figure 7.**
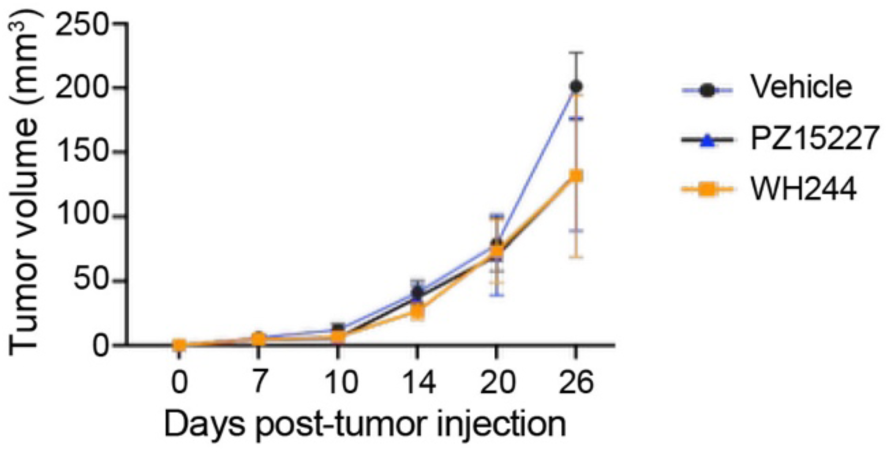
LVRCC67 tumors are resistant to single therapy using Bcl-X_L_-targeting PROTAC. Balb/C mice bearing Renca tumors were treated with vehicle, PZ15227, or WH244 (n=5 per group). Tumor volume was recorded and graphed (n=5 per group).

## Notes

### Competing Interest Statement

Conflict of Interest: G.Z. and W.Z. are equity holders of Dialectic Therapeutics, a company that is developing BCL-XL/BCL-2 PROTACs to treat cancers; G.Z., W.Z., P.Z. have also several patents for developing BCL-XL/BCL-2 PROTACs to treat cancers.

